# The origins of haplotype 58 (H58) *Salmonella enterica* serovar Typhi

**DOI:** 10.1101/2022.10.03.510628

**Authors:** Megan E. Carey, To Nguyen Thi Nguyen, Tran Do Hoang Nhu, Zoe A. Dyson, Pham Thanh Duy, Elli Mylona, Satheesh Nair, Marie Chattaway, Stephen Baker

**Affiliations:** Cambridge Institute of Therapeutic Immunology & Infectious Disease (CITIID), Department of Medicine, University of Cambridge, Cambridge, UK; The Hospital for Tropical Diseases, Wellcome Trust Major Overseas Program, Oxford University Clinical Research Unit, Ho Chi Minh City, Vietnam; Department of Microbiology, Monash Biomedicine Discovery Institute, Monash University, Melbourne, Victoria, Australia; School of Medicine, Stanford University, Stanford, USA; London School of Hygiene & Tropical Medicine, London, UK; Wellcome Sanger Institute, Wellcome Genome Campus, Hinxton, Cambridge, UK; United Kingdom Health Security Agency, Gastrointestinal Bacteria Reference Unit, London, UK

**Keywords:** *Salmonella* Typhi (*S*. Typhi), antimicrobial resistance, H58, chronic carriage, genomics

## Abstract

Antimicrobial resistance (AMR) poses a serious threat to the clinical management of typhoid fever. AMR in *Salmonella* Typhi (*S*. Typhi) is associated with the H58 lineage, which arose comparatively recently before becoming globally disseminated. To better understand when and how this lineage emerged and became dominant, we performed detailed phylogenetic and phylodynamic analyses on contemporary genome sequences from *S*. Typhi isolated in the period spanning the emergence. Our dataset, which contains the earliest described H58 *S*. Typhi, indicates that the prototype H58 organisms were multi-drug resistant (MDR). These organisms emerged spontaneously in India in 1987 and became radially distributed throughout South Asia and then globally in the ensuing years. These early organisms were associated with a single long branch, possessing mutations associated with increased bile tolerance, suggesting that the first H58 organism was generated during chronic carriage. The subsequent use of fluoroquinolones led to several independent mutations in *gyrA*. The ability of H58 to acquire and maintain AMR genes continues to pose a threat, as extensively drug-resistant (XDR; MDR plus resistance to ciprofloxacin and third generation cephalosporins) variants, have emerged recently in this lineage. Understanding where and how H58 *S*. Typhi originated and became successful is key to understand how AMR drives successful lineages of bacterial pathogens. Additionally, these data can inform optimal targeting of typhoid conjugate vaccines (TCVs) for reducing the potential for emergence and the impact of new drug-resistant variants. Emphasis should also be placed upon the prospective identification and treatment of chronic carriers to prevent the emergence of new drug resistant variants with the ability to spread efficiently.

## Introduction

*Salmonella enterica* serovar Typhi (*S*. Typhi) is the etiologic agent of typhoid fever, a disease associated with an estimated 10.9 million new infections and 116,800 deaths annually.^1^ The disease classically presents as a non-differentiated fever and can progress to more severe manifestations or even death.^2^ Typhoid fever necessitates antimicrobial therapy, as the associated mortality rate in the pre-antimicrobial era ranged from 10–30%;^3^ presently, typhoid has a case fatality rate (CFR) of <1% when treated with effective antimicrobials.^4^ *S*. Typhi is spread via the faecal-oral route, typically through the ingestion of contaminated food or water.^2^ Therefore, high prevalence rates of typhoid fever were historically associated with urban slums in South Asia with poor sanitation.^5^ Recent multicentre surveillance studies have demonstrated that typhoid fever is also a major problem in both urban and rural areas in sub-Saharan Africa.^6–8^

Given the importance of antimicrobials for the management and control of typhoid, antimicrobial resistance (AMR) in *S*. Typhi has the potential to be a major public health issue. Indeed, the problem of AMR in *S*. Typhi first appeared in the 1950s with the emergence of resistance against the most widely used drug, chloramphenicol.^9^ Multi-drug resistant typhoid (MDR; resistance to all first-line antimicrobials chloramphenicol, trimethoprim-sulfamethoxazole, and ampicillin) was first identified in the 1970s and became common in the early 1990s.^10,11^ MDR in *S*. Typhi is frequently conferred by self-transmissible IncH1 plasmids carrying a suite of resistance genes, include resistance determinants for chloramphenicol (*catA1* or *cmlA*), ampicillin (*bla*TEM-1D, *bla*OXA-7), and co-trimoxazole (at least one *dfr*A gene and at least one *sul* gene).^12^ Lower efficacy of first-line antimicrobials led to the increased use of fluoroquinolones, but decreased fluoroquinolone susceptibility became apparent in the mid-1990s, and was widespread in South and Southeast Asia in the early 2000s.^13,14^ Inevitably, as treatment options have become limited, third-generation cephalosporins and azithromycin have been used more widely for effective treatment of typhoid fever.^15–17^ However, newly circulating extensively-drug resistant variants of *S*. Typhi (XDR; MDR plus resistance to fluoroquinolones and third generation cephalosporins) has left azithromycin as the only feasible oral antimicrobial for the treatment of typhoid fever across South Asia.^18^ We are arguably at a tipping point, as azithromycin-resistant *S*. Typhi has since been reported in Bangladesh, Pakistan, Nepal, and India, thereby threatening efficacy of common oral antimicrobials for effective typhoid treatment.^19–22^ If an XDR organism were to acquire azithromycin resistance (single bae pair mutation), this would lead to what Hooda and colleagues have referred to as pan-oral drug-resistant (PoDR) *S*. Typhi, which would require inpatient intravenous treatment.^23^ This would come at substantial additional cost to patients and their families, and place additional strain on already overburdened health systems. ^24–26^

In contrast to many other Gram-negative bacteria, *S*. Typhi is human restricted with limited genetic diversity that can be described by a comparatively straightforward phylogenetic structure.^27^ Therefore, the phylogeny and evolution of *S*. Typhi provide a model for how AMR emerges, spreads, and becomes maintained in a human pathogen. AMR phenotypes in *S*. Typhi are typically dominated by a single lineage; H58 (genotype 4.3.1 and consequent sublineages), which was the 58^th^ *S*. Typhi haplotype to be described in the original genome wide typing system.^28^ This highly successful lineage is commonly associated with MDR phenotypes and decreased fluoroquinolone susceptibility.^14^ Previous phylogeographic analysis suggested that H58 emerged initially in Asia between 1985 and 1992 and then disseminated rapidly to become the dominant clade in Asia and subsequently in East Africa.^14^ H58 is currently subdivided into three distinct lineages – lineage I (4.3.1.1) and lineage II (4.3.1.2), which were first identified in a pediatric study conducted in Kathmandu,^29^ and lineage III (4.3.1.3), which was identified in Dhaka, Bangladesh.^30^ A recent study of acute typhoid fever patients and asymptomatic carriers in Kenya demonstrated the co-circulation of genotypes 4.3.1.1 and 4.3.1.2 in this setting, and closer analysis showed that these East African sequences had distinct AMR profiles and were the result of several introduction events.^31,32^ These events led to the designation of three additional genotypes: H58 lineage I sublineage East Africa I (4.3.1.1.EA1), H58 lineage II sublineage East Africa II (4.3.1.2.EA2), and H58 lineage II sublineage East Africa III (4.3.1.2.EA3).^32^ In addition, the XDR *S*. Typhi clone, which was caused by a monophyletic outbreak of genotype 4.3.1.1 organisms, was designated genotype 4.3.1.1.P1 to facilitate monitoring of its spread.

It is apparent from investigating the phylogeny of *S*. Typhi that H58 is atypical in comparison to other lineages. This lineage became dominant in under a decade, and first appeared on a long basal branch length, indicative of a larger number of single base pair mutations separating it from its nearest neighbour (Figure 1a). These observations suggest that there is something ‘unique’ about the evolution of this lineage, but we have limited understanding of how H58 emerged, what enabled its rapid spread, and when it initially appeared. Here, by collating new genome sequences of *S*. Typhi that were associated with travel to South Asia in the late 1980s and early 1990s and comparing them to a global population over the same period, we explore an expanded early phylogenetic dataset to resolve the origins and rapid success of this important and successful AMR clone.

**Figure 1.**
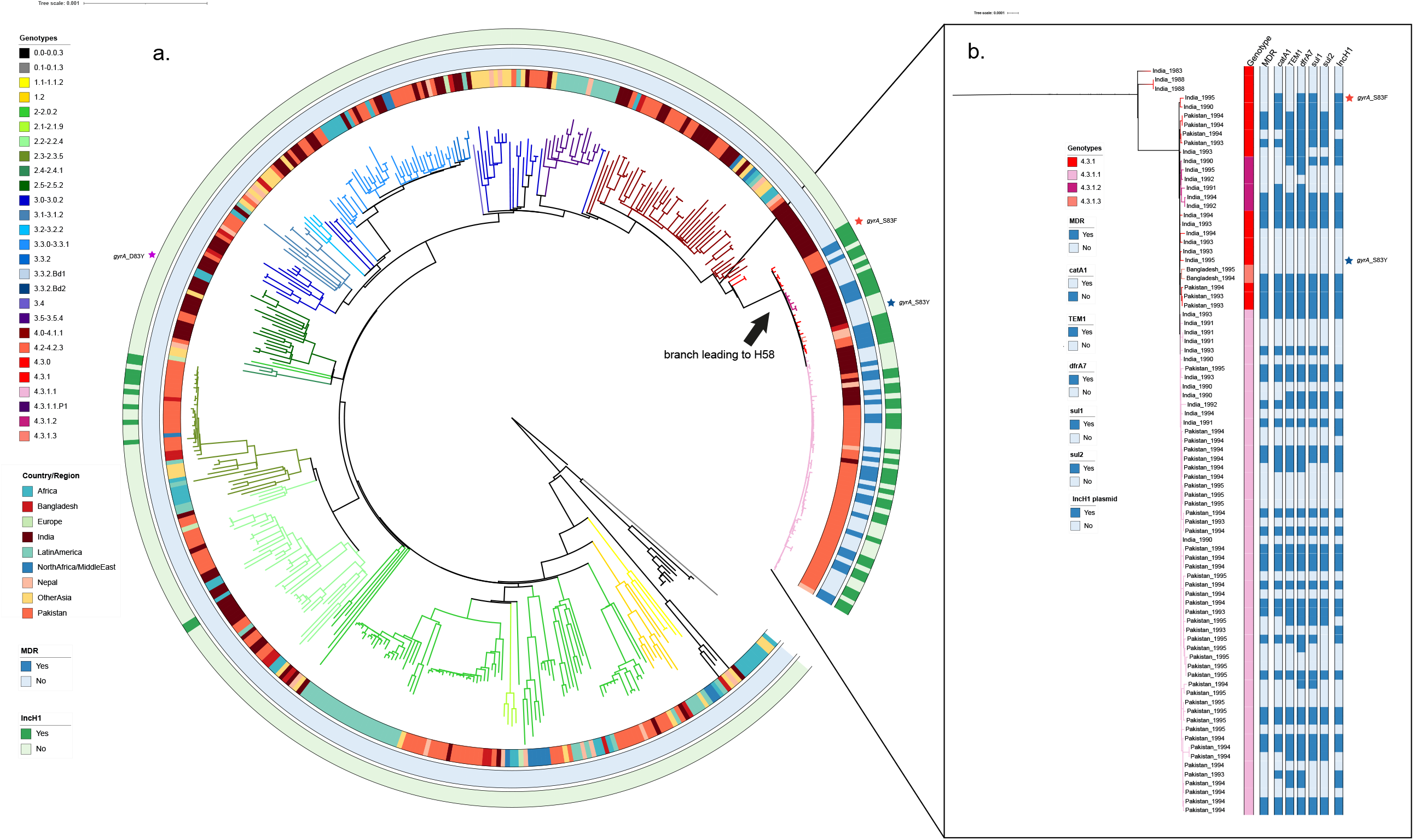
The phylogenetic structure of historical *S*. Typhi isolates. a) A phylogenetic overview of historical (1980 – 1995) *S*. Typhi from the UKHSA. Maximum likelihood outgroup rooted phylogenetic tree depicting the genomic sequences of 463 *S*. Typhi isolated from returning travellers to the United Kingdom isolated between 1980 and 1995. Branch colour indicates genotype, and rings outside of the tree indicate country of origin, MDR, and presence of IncH1 plasmid, coloured as per the inset legend. Individual mutations in the quinolone resistance determining region (QRDR) indicated by stars outside the rings. b) A phylogenetic overview of historical H58 (4.3.1) *S*. Typhi isolates. Zoomed-in view of H58 *S*. Typhi isolates from historical collection. Genotype, country of origin, presence of MDR, AMR mutations, and presence of IncH1 plasmids are indicated by bars to the right of the tree and coloured as per the inset legend. Individual mutations in the quinolone resistance determining region (QRDR) indicated by stars to the right of the bars.

## Results and discussion

### Sampling

The main questions that we aimed to address with this study were: i) when and where did H58 *S*. Typhi first emerge; ii) can we better resolve the evolutionary events that lead to the long branch length observed for H58 *S*. Typhi; and iii) how quickly did this lineage spread and why? Therefore, to investigate the origins of *S*. Typhi H58, data from United Kingdom Health Security Agency (UKHSA, formerly Public Health England) containing information on stored *S*. Typhi organisms isolated between 1980 and 1995 from travellers returning to the UK from overseas and receiving a blood culture were analysed.

The database was queried and organisms were selected from the following three categories: i) 126 *S*. Typhi with the E1 Phage type (which is considered to be associated with H58)^12^ originating from South Asia (India, Nepal, Pakistan, and Bangladesh), ii) 159 *S*. Typhi organisms with a variety of non-E1 phage types originating from South Asia, and iii) 184 *S*. Typhi organisms with a variety of phage types (both E1 and non-E1) originating from locations outside of South Asia. A total of 470 *S*. Typhi organisms meeting these criteria were randomly selected, revived, subjected to DNA extraction and whole genome sequenced. Ultimately, our dataset was composed of 463 novel sequences generated as a component of this study and 305 existing sequences^30,33,34^ known to belong to the H58 lineage and its nearest neighbours, yielding a total of 768 whole genome sequences on which to structure subsequent analysis.

### Population structure, genotype distribution, and antimicrobial resistance profiles of historical S. Typhi

We inferred a maximum likelihood phylogenetic tree from the sequencing data to examine the population structure of this historic (1980 – 1995) collection of global *S*. Typhi sequences (Figure 1). Notably, unlike the extent global population of *S*. Typhi, which is largely dominated by a single lineage,^14,35^ this historic population exhibited considerable genetic diversity, with 37 genotypes represented (Figure 1a). The majority of isolates belonged to primary clade 2 (194/463), of which clade 2.0 was most common (36%, 69/194) followed by subclades 2.3.3 (14%, 28/194) and 2.2.2 (13%, 25/194). An additional 23% of isolates belonged to primary clade 3 (108/463) and 3% of isolates were classified as major lineage 1 (12/463). Ultimately, 29% of isolates belonged to major lineage 4 (134/463), of which 63% (84/134) were H58. Among these, H58 lineage I (genotype 4.3.1.1) was most common (67%, 56/84), followed by genotype 4.3.1 (H58 not differentiated into any sublineage; 24%, 20/84). The earliest H58 isolates in our dataset are illustrated in higher resolution in Figure 1b.

Although we enriched our historical dataset for samples isolated from travellers returning from South Asia, our final dataset generated substantial geographic coverage, with 39 different countries represented. Therefore, these data are likely representative of the circulating *S*. Typhi in this pivotal period. Notably, the earliest H58 organism in our dataset that was classified as H58 according to the GenoTyphi scheme was isolated in 1983 from an individual entering the UK from India, followed by two additional Indian isolates (1988) that were also classified as H58. These organisms differed from the larger cluster of H58 organisms by 61-64 single nucleotide polymorphisms (SNPs) (Figure 1a). All H58 *S*. Typhi in this dataset were isolated from travellers returning from South Asia, with the majority (51/84; 61%) originating from Pakistan and the remainder from India and Bangladesh; 31/84 (37%) and 2/84 (2%), respectively.

Given that this dataset included isolates from the early MDR era and then following the emergence of reduced fluoroquinolone susceptibility, we analysed the data for genes associated with MDR and mutations in the DNA gyrase gene, *gyrA*. Overall, 7% (34/463) of the organisms in this historical dataset were genetically defined as being MDR; significantly, all were H58 (genotypes 4.3.1, 4.3.1.1, 4.3.1.2, and 4.3.1.3) and isolated between 1991 and 1995, and 97% (33/34) possessed an IncH1 plasmid (Figure 1b). Thirteen H58 organisms contained an IncH1 plasmid carrying AMR genes, but did not possess the genes conferring resistance to all three first-line antimicrobials, and thus were not genetically defined as MDR (Figure 1b, Table S1). The first mutation in the quinolone resistance determining region (QRDR) in our dataset was a *gyrA*-D87Y substitutions identified in a genotype 2.5 organism originating in South Africa in 1986 (Figure 1a). This was clearly a spontaneous mutation that evolved *de novo* and did not appear to become fixed in the population. Similarly, two single QRDR mutations (*gyrA*-S83F and *gyrA*-S83Y) occurred independently in H58 organisms (genotype 4.3.1) in India in 1995 (Figure 1b) and were not observed in descendant populations. No mutations in *gyrB* or *parC* were observed in this dataset.

In order to contextualise these isolates to understand the evolutionary events leading to this clone, we selected H58 and nearest neighbours (from genotypes 4.1 and 4.2) *S*. Typhi organisms (n=305) that were already available in the public domain from previous studies^30,33,34^ (Table S2) and generated a phylogenetic tree combining these isolates with early H58 and nearest neighbour isolates from our unpublished dataset (n=117). In our H58 and nearest neighbour dataset (n= 422, Figure 2), which included both published data as well as our contemporary data, 17 countries were represented.^30,33,34^ Of the non-H58 isolates (nearest neighbours), 42% were genotype 4.1 (32/76), 13% (10/76) were genotype 4.2, 28% (21/76) were 4.2.1, and 16% (12/76) were 4.2.3. These non-H58 nearest neighbour organisms were isolated between 1981 and 2000. None of them were MDR, and none carried an IncH1 plasmid. Of our H58 isolates (defined as having informative SNPs indicative of lineage 4.3.1), the earliest organism was isolated in 1983 in India, followed by two additional Indian isolates (1988) that were also classified as H58. However, we can observe that there were no more recent isolates from this founder group (Figure 2), implying that this lineage became extinct; these isolates did not contain incH1 plasmids and were non-MDR. Within the H58 lineage, most of the organisms belonged to H58 sublineage I (4.3.1.1; 84%, 290/346), followed by sublineage II (4.3.1.2; 7%, 25/346), genotype 4.3.1 (6%, 21/346), and sublineage III (4.3.1.3; 3%, 10/346). Overall, 63% (219/346) of these H58 organisms were MDR, and 87% (191/219) of these MDR H58 organisms carried an IncH1 plasmid. All of the MDR H58 organisms lacking an IncH1 plasmid were genotype 4.3.1.1, the earliest of which was isolated in India in 1991.Within this group, the first single point mutation in the QRDR occurred comparatively early in an organism isolated in India in 1991.^33^

**Figure 2.**
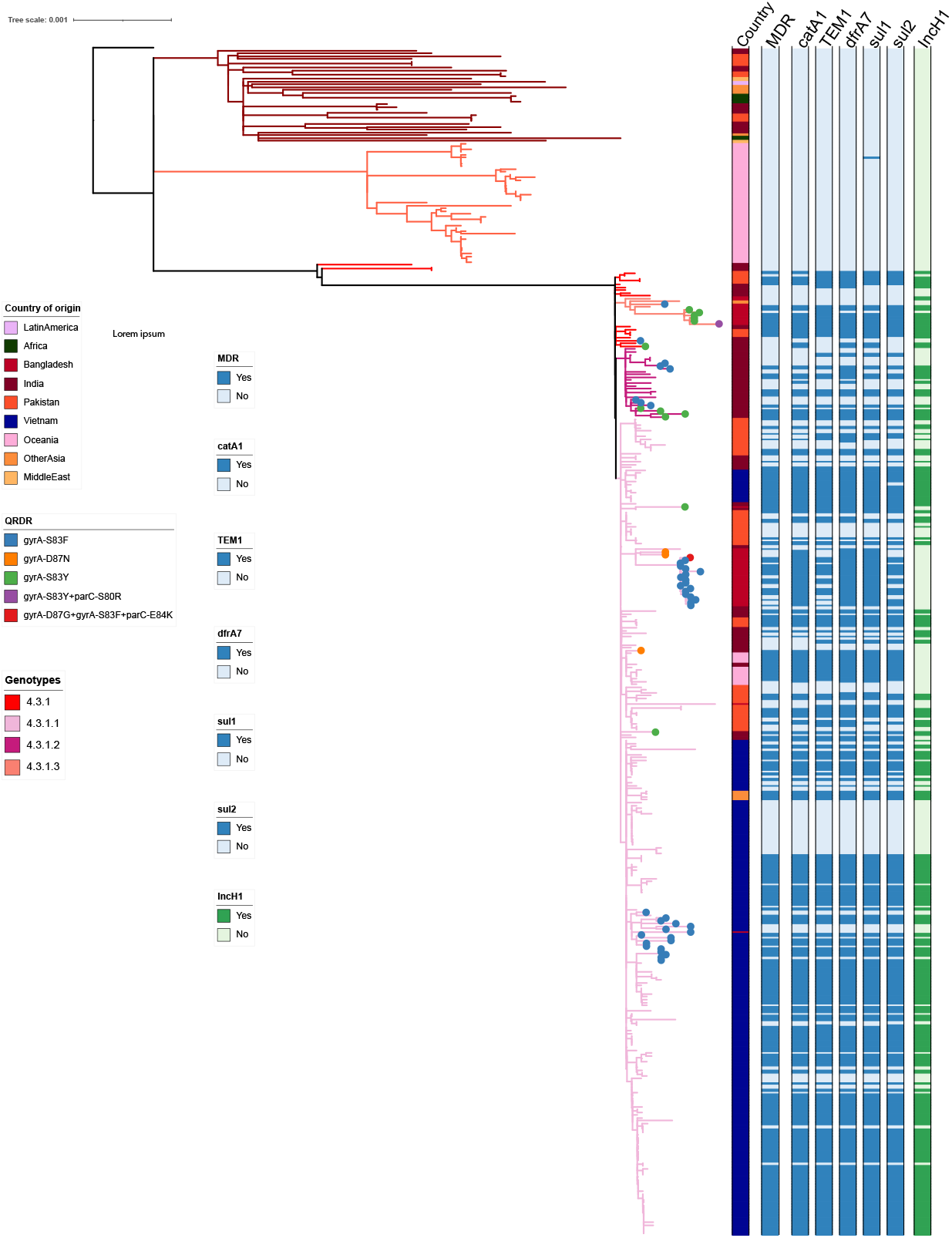
The phylogenetic structure of early H58 organisms and nearest neighbours. Maximum likelihood rooted phylogenetic tree showing *S*. Typhi organisms (genotype 4.3.1) and nearest neighbours (genotypes 4.1 and 4.2) from our historical collection and from published literature (n=422 total). Genotype is indicated by branch colour, presence of QRDR mutation(s) are indicated by coloured circles at the end of the branches, and country of origin, presence of MDR, AMR mutations, and presence of incH1 plasmid are indicated by bars to the right of the tree and coloured as per the inset legend.

### Evolutionary history of H58 S. Typhi

Using BEAST analysis, we determined that the median substitution rate of the H58 was 2.79 × 10^−7^ substitutions base^−1^ year^−1^ [95% highest posterior density (HPD): 2.40 × 10^−7^ -3.24 × 10^−7^), which is comparable to that observed in previous studies.^36^ We found that the most recent common ancestor (MRCA) of the H58 was estimated to have emerged in late 1987 (95% HPD: 1986 – 1988). Two H58 sublineages (4.3.1.1 and 4.3.1.2) then emerged almost simultaneously in India in 1987 and 1988. The time-inferred phylogeny shows a clonal expansion of H58 that originated from South Asia, specifically in India, and then disseminated globally. As noted above, 63% (219/346) of isolates were MDR, and 87% (191/219) of those isolates contained an IncH1 plasmid known to carry AMR genes. Detailed genetic analysis of the IncH1 plasmids observed in most of these MDR isolates revealed high genetic similarity, with only 13 single nucleotide polymorphisms (SNPs) difference between them (Supplementary Figure 1). These data strongly suggest that the ancestral H58 organism that was the basis for the major clonal expansion was already MDR before undergoing clonal expansion and subsequent global dissemination; some ensuing H58 organisms then lost the MDR plasmid in certain settings, presumably because of decreased antimicrobial selection pressure with first-line antimicrobials, with a corresponding impact on fitness. The fact the three early precursor organisms that were not MDR appear to have become extinct supports our hypothesis that the presence of an MDR phenotype was a pivotal selective event.

It is also stark that QRDR mutations appeared in H58 organisms quickly and frequently. Within this H58 dataset, organisms with one or more QRDR mutations appeared within six independent lineages of H58, as illustrated in Figures 2 and 3. The earliest H58 lineage (lineage II) to develop QRDR mutations was observed in 1991 in India, with a single S83Y mutation in the *gyrA* gene, and the first isolate containing two QRDR mutations (*gyr*AS83Y; *par*C-S80R) appeared in a genotype 4.3.1.3 organism in Bangladesh in 1999 (Figure 3). The first “triple mutant” (mutations in *gyr*A-D87G, *gyr*A-S83F, and *parC*-E84K) was found in a 4.3.1.1 Bangladeshi isolate in 1999 (Figure 3).

**Figure 3.**
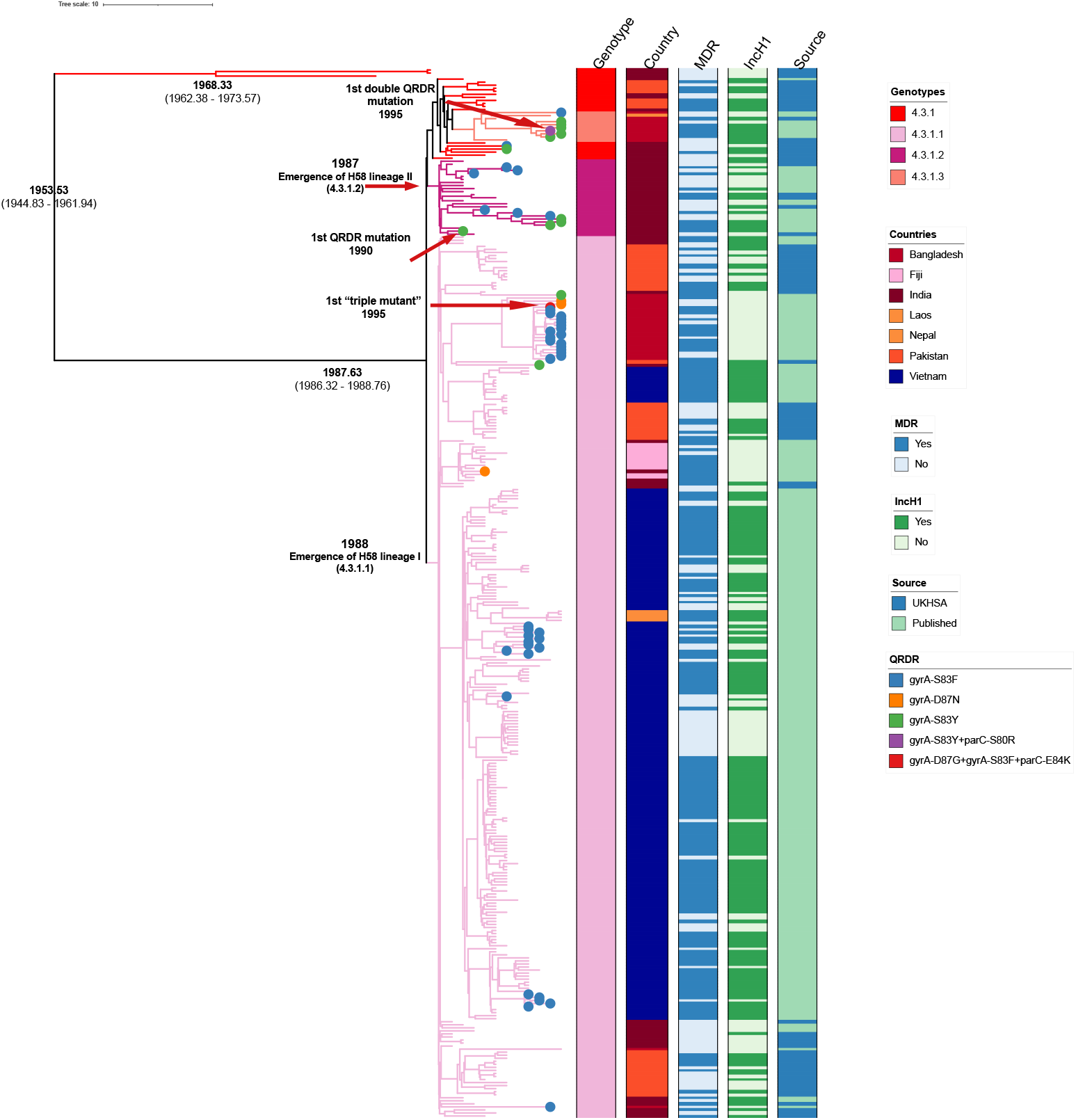
A dated phylogenetic structure of historical H58 *S*. Typhi isolates. BEAST-generated dated phylogeny of H58 *S*. Typhi isolates from UKHSA collection and published literature (n=345). Tip colours indicate presence of specific mutation(s) in the QRDR as per inset legend. Branch colour and the first column to the right of the tree indicate genotype, the second column indicates country of origin, the third represents presence of MDR, and the final column indicates presence of an incH1 plasmid. Our analysis suggests that the Most Recent Common Ancestor (MCRA) of H58 appeared in 1987, and that two sublineages (I and II) emerged almost simultaneously in India in 1987 and 1988. The first single point mutation in the QRDR was observed in 1990, and the first “triple mutant” was observed in Bangladesh in 1999.

### Genetic variation associated with H58 S. Typhi

Our data support the hypothesis that H58 *S*. Typhi was successful specifically because of the acquisition and maintenance of an MDR plasmid. This selection meant that later *gyrA* mutations were more likely to occur in this lineage given its dominance (and therefore, higher rates of replication leading to additional opportunities for mutations to occur), as well as assumed frequent fluoroquinolone exposure, given its existing MDR phenotype. Based on the results of our mapping, we undertook further genetic analysis to identify non-synonymous SNPs unique to the early H58 isolates, as well as SNPs that were unique to early H58 isolates that were MDR. The motivation was to explain the origins of the long branch length illustrated on Figure 1a, to infer why this lineage was so globally successful, and identify genetic elements that may stabilize an MDR IncH1 plasmid.

We identified 16 unique non-synonymous SNPs that were exclusive to the early H58 isolates as compared to precursor 4.1 and 4.2 organisms, the majority of which were present in genes associated with central metabolism and outer membrane structures; one of which was associated with pathogenicity (Table 1). Within the early H58 isolates that were also MDR, we identified an additional 23 unique non-synonymous SNPs, most of which were found in genes encoding proteins predicted to regulate metabolism, degrade small molecules, membrane/surface structures, as well as regulators, pathogenicity adaptation, and information transfer (Table 2). We additionally identified mutations in a gene (t2518/STY0376) encoding a hypothetical protein with an EAL (diguanylate phosphodiesterase) domain with a non-synonymous mutation, which has previously been identified as being associated with H58 organisms.^37^ The homologous gene (STM0343) in *Salmonella* Typhimurium has been described as regulating motility and invasion, which suggests that SNPs in this gene might also contribute to virulence.^38,39^ In addition, we identified SNPs in genes that have previously been associated with tolerance to bile in *S*. Typhi (*sirA, recB, wecF, dsdA*, and *yjjV*) among the early MDR H58 isolates.^40,41^

**Table 1.** Unique non-synonymous Single Nucleotide Polymorphisms detected in early H58 *S*. Typhi

| Gene ID | SNP Position in CT18 | Gene | Reference nucleotide | 4.3.1 nucleotide | Ancestral codon | Derived codon | Ancestral amino acid | Derived amino acid | Functional Category | Product |
| --- | --- | --- | --- | --- | --- | --- | --- | --- | --- | --- |
| STY0376 | 387595 |  | C | T | ACC | ATC | T | I | Central/intermediary metabolism | putative rtn protein; diguanylate cyclase/ phosphodiesterase domain-containing protein |
| STY0452 | 461438 | *yajI* | G | A | CGT | TGT | R | C | Membrane/surface structures | putative lipoprotein |
| STY0522 | 529155 | *kefA/aefA* | A | G | AAA | GAA | K | E | Membrane/surface structures | integral membrane protein AefA |
| STY0698 | 693560 | *rlpB* | C | T | ATG | ATA | M | I | Central/intermediary metabolism | rare lipoprotein B precursor |
| STY1458 | 1408039 |  | C | T | CAG | TAG | Q | * |  | putative lipoprotein |
| STY1703 | 1629304 | *ssaP* | G | A | GCG | GTG | A | V | Pathogenicity/adaptation/c haperones | putative type III secretion protein |
| STY2371 | 2202853 |  | C | T | CGT | TGT | R | C | Membrane/surface structures | putative nucleoside permease |
| STY2513 | 2348633 | *glpA* | G | A | GGC | AGC | G | S | Central/intermediary metabolism | anaerobic glycerol-3-phosphate dehydrogenase subunit A |
| STY2553 | 2388057 | *nuoG* | G | A | ACT | ATT | T | I | Information transfer | NADH dehydrogenase I chain G |
| STY2564 | 2401233 | *yfbT* | G | A | CGC | TGC | R | C | Central/intermediary metabolism | putative phosphatase |
| STY3518 | 3360344 | *nanE2* | T | C | ATA | GTA | I | V | Pseudogenes | conserved hypothetical protein (pseudogene) |
| STY3795 | 3659647 | *Ydey* | C | T | GGG | GAG | G | E | Membrane/surface structures | Putative ABC transporter protein, AI-2 transport system permease |
| STY4405 | 4273783 | *metH* | C | A | CGT | AGT | R | S | Central/intermediary metabolism | B12-dependent homocysteine-N5-methyltetrahydrofolate transmethylase |
| STY4451 | 4321540 |  | A | G | AAG | AGG | K | R |  | single-strand DNA-binding protein |
| STY4890 | 4754034 |  | C | T | TGG | TAG | W | * |  | probable carbon starvation protein |
| STY4890 | 4754035 |  | A | G | TGG | CGG | W | R |  | probable carbon starvation protein |

**Table 2.** Unique non-synonymous SNPs unique to early MDR H58 *S*. Typhi

| Gene ID | SNP Position in CT18 | Gene | Reference nucleotide | 4.3.1 nucleotide | Ancestral codon | Derived codon | Ancestral amino acid | Derived amino acid | Functional Category | Product |
| --- | --- | --- | --- | --- | --- | --- | --- | --- | --- | --- |
| STY0042 | 40159 | *betC* | G | A | GGG | GAG | G | E | Central/intermediary metabolism | putative secreted sulfatase |
| STY0085 | 89102 | *etfB/fixA* | A | G | AAA | AGA | K | R | Degradation of small molecules | FixA protein |
| STY0376 | 387082 |  | G | A | CGA | CAA | R | Q | Central/intermediary metabolism | putative rtn protein; diguanylate cyclase/phosphodiesterase domain-containing protein |
| STY0886 | 880083 | *ybiK* | A | G | ACC | GCC | T | A | Central/intermediary metabolism | putative L-asparaginase |
| STY1310 | 1270888 |  | G | A | CGA | CAA | R | Q | Membrane/surface structures | Voltage-gated potassium channel |
| STY1328 | 1286044 | *trpE* | T | C | GAC | GGC | D | G | Central/intermediary metabolism | Anthranilate synthase component |
| STY1410 | 1360939 | *dbpA* | C | T | CAG | TAG | Q | * | Pseudogenes | ATP-dependent RNA helicase (pseudogene) |
| STY1917 | 1810914 | *hyaE* | G | A | GCT | ACT | A | T | Information transfer | hydrogenase-1 operon protein HyaE |
| STY2155 | 2002943 | *uvrY* | G | A | CTT | TTT | L | F | Regulators | invasion response-regulator |
| STY2875 | 2750755 |  | G | A | GCG | ACG | A | T | Pathogenicity adaptation/chaperones | large repetitive protein |
| STY3001 | 2875160 | *sptP* | G | A | CAA | TAA | Q | * | Pathogenicity adaptation/chaperones | Pathogenicity island 1 tyrosine phosphatase (associated with virulence) |
| STY3132 | 3004181 | *recB* | T | C | GAA | GGA | E | G | Degradation of macromolecules | exonuclease V subunit |
| STY3297 | 3144053 | *ordL* | A | C | GTG | GGG | V | G | Central/intermediary metabolism | Putative gamma-glutamylputrescine oxidoreductase |
| STY3555 | 3398551 | *yhdA* | G | A | GCT | GTT | A | V | Membrane/surface structures | Putative lipoprotein |
| STY3628 | 3484294 | *wecF* | C | T | GTA | ATA | V | I | Conserved hypothetical proteins | Putative 4-alpha-L-fucosyl transferase |
| STY3955 | 3824631 | *torC* | T | G | TCA | GCA | S | A | Pseudogenes | Cytochrome c-type protein |
| STY3977 | 3843665 | *dsdA* | C | T | GGC | GAC | G | D | Degradation of small molecules | D-serine dehydratase |
| STY4161 | 4020211 | *yhjY* | C | T | ACG | ATG | T | M | Membrane/surface structures | putative membrane protein |
| STY4314 | 4192687 | *gph* | C | T | GCG | GTG | A | V | Degradation of small molecules | phosphoglycolate phosphatase |
| STY4318 | 4196909 | *bigA* | G | A | CCA | TCA | P | S | Pseudogenes | putative surface-exposed virulence protein (pseudogene) |
| STY4392 | 4253640 | *dprA* | G | A | GCT | ACT | A | T | Conserved hypothetical proteins | Putative DNA protecting protein |
| STY4805 | 4665891 |  | G | A | GCG | GTG | A | V | Regulators | arginine deiminase |
| STY4915 | 4775254 | *yjjV* | C | A | CCG | CAG | P | Q | Conserved hypothetical proteins | TatD DNase family protein |

Clearly, there was a cascade of events that corresponded with the genesis of the first H58 *S*. Typhi. The cumulative mutations signified by the observed long branch length is uncommon in *S*. Typhi and has two feasible explanations. The first is that the progenitor organism was a hyper mutator, and that a key mutation in *mutS* was responsible for generating a large amount of genetic diversity in a short time frame.^42^ However, no such informative SNPs were observed in the early H58 isolates, but any mutations may have reverted. The second, and more likely explanation, is that the organism was in an environment that created an atypical selective pressure to induce mutations that facilitated its ability become exposed to, and then accept, an MDR plasmid. Our previous data on *S*. Typhi carriage in the gallbladder determined that this environment creates an atypical selective pressure and stimulates mutations in metabolism and outer membrane structures.^31,43^ This genetic variation was associated with organisms being located on signature long branches; our observations here are comparable. We suggest that H58 *S*. Typhi became successful due to its early ability to accept and stabilise a large MDR plasmid, which probably occurred whilst in the gallbladder; this one-off event and onward transmission then created this successful lineage. Therefore, we speculate that gallbladder carriage acts as a niche for generation of new variants with both modest (single SNPs) and large (plasmid acquisition) events capable of generating new lineages of *S*. Typhi with a selective advantage.

### Implications

Our phylodynamic analysis of the origins of H58 *S*. Typhi has important consequences for how we understand the emergence and spread of new drug-resistant variants and can help inform optimal use of typhoid conjugate vaccines (TCVs). Our data suggest that whilst rare, these events can happen given the specific selective pressures, allowing MDR organisms to arise and spread rapidly. This observation comes at a critical time in global typhoid control. Two TCVs have been prequalified by the World Health Organization, with additional candidates in late-stage clinical development, and promising clinical efficacy and effectiveness data.^44,45^ The continued spread of drug-resistant H58 genotypes is clearly a major argument for the use of TCV, particularly as resistance to all oral antimicrobials has been reported in *S*. Typhi in South Asia.^19–22^ We continue to observe a pattern in which drug-resistant variants emerge in South Asia and spread radially, with H58 being the only major genotype to do so to date. ^14,35,46–49^ H58 *S*. Typhi was first isolated in Kenya shortly after its estimated emergence in 1988, further illustrating the potential for rapid spread of this lineage.^48^ This suggests that prioritization of widespread use of TCVs in South Asia can not only prevent significant morbidity in the region but should limit the continued emergence and spread of drug-resistant organisms elsewhere, which could expand the useful lifespan of existing therapeutic options in some parts of the world until TCVs become more widely available.

Our data strongly suggest that H58 *S*. Typhi, which is highly associated with being MDR and decreased fluoroquinolone susceptibility, is highly adept at acquiring and maintaining drug resistance determinants, which is likely facilitated by unique mutations that occurred in the earliest H58 organism. The rapid international dissemination of the H58 lineage, starting in South Asia and spreading throughout Southeast Asia,^14,33^ into Africa^14,31,47,48,50–52^ and more recently, Latin America,^53^ suggest that expanded genomic surveillance is warranted to monitor its continued global spread. Such information can also help inform the development of transmission dynamics models that predict the spread of newer drug-resistant variants, like XDR, which can also inform TCV introduction decision-making.

Ultimately, we surmise that H58 *S*. Typhi likely emerged in India from a chronic carrier, which is supported by the indicative long branch length between early H58 organisms and its nearest non-H58 neighbours. This deduction is consistent with observations from previous studies conducted in Nepal and Kenya, in which higher mean branch lengths were observed in carriage isolates as compared to isolates from symptomatic patients;^31,43^ this phenomenon is to be expected, assuming that chronic carriers will have had a longer time from acquisition of infection to shedding and sampling. Notably, considering the structure of the phylogenetic tree and the loss of the early non-MDR H58 organisms from the population, it is likely that the MDR phenotype was the main catalysing factor for the success of this lineage. Additionally, we observed non-synonymous mutations in genes associated with outer membrane structures, metabolism and virulence, which we have been observed previously in organisms isolated directly from the gallbladder.^31,43^ Our analysis also supports previous transcriptomic analysis showing that H58 *S*. Typhi has higher bile tolerance relative to other laboratory strains (Ty2 and CT18) and show increased virulence in the presence of bile, thereby increasing the potential of H58 organisms to colonize and persist in the gallbladder. These observations suggest that gallbladder is the ideal location for the generation of variants and highlights the potential for chronic carriage to lead to the emergence of novel *S*. Typhi (and other invasive *Salmonella*) that are genetically predisposed to express new phenotypes, which may include drug resistance. Therefore, emphasis should be placed upon the prospective identification and treatment of chronic carriers to prevent the emergence of new variants with the ability to spread. Contemporary data from returning travellers to the UK suggest that 1.4% of those infected with *S*. Typhi are chronic carriers, and 0.7% are carrying MDR *S*. Typhi.^54^ A comparable frequency of carriage (1.1%) has been observed among children aged 16 years and younger in Mukuru, an informal settlement close to Nairobi, Kenya.^31^ This prevalence rate is likely to be higher in older age groups,^55–57^ and among people living in settings where typhoid is hyperendemic, and thus, may present a more substantial risk in terms of sustained transmission of drug-resistant *S*. Typhi and the potential emergence of additional drug-resistant variants. Scalable, low-cost assays to detect carriers will become vital if we aim to eliminate typhoid and prevent future resurgence.

We conclude that H58 *S*. Typhi likely emerged from a chronic carrier in India in 1987. The prototype organism of the successful clonal expansion was already MDR and became highly successful across South Asia in over a period of <10 years. Ultimately, sustained use of, and exposure to, fluoroquinolones led to selective mutations in *gyrA* on many independent occasions. The dominance of this organism and its ability to maintain AMR genes has latterly meant it has become resistant to additional antimicrobials. Our work represents a blueprint of how such organisms can arise and become dominant, but also provides the justification and evidence for the introduction of new interventions for disease control; if we reduce disease burden by vaccination, we will additionally reduce the likelihood of comparable events occurring in other *S*. Typhi organisms and other pathogens. Widespread vaccine deployment, as well as screen and treat programs, can not only impact AMR directly through the prevention of drug-resistant infections, but also indirectly, as reduced transmission leads to decreased selection pressure on account of lower bacterial replication, and potentially decreased antimicrobial use following the prevention of clinical disease warranting treatment.

## Methods

### DNA extraction and Whole Genome Sequencing

Genomic DNA from *S*. Typhi isolates was extracted using the Wizard Genomic DNA Extraction Kit (Promega, Wisconsin, USA), following standardized manufacturer’s protocol. Two ng of genomic DNA from each organism was fragmented and tagged for multiplexing with Nextera DNA Sample Preparation Kits, followed by paired-end sequencing on an Illumina HiSeq2000 Platform to produce 101 bp paired-end reads (Illumina, Cambridge, UK). Raw reads were deposited in the European Nucleotide Archive (ENA) under study accession number PRJEB15284 (Table S1).

### Read alignment and SNP analysis

FastQC and FASTX-Toolkit bioinformatics pipelines were used to check the quality of raw reads.^58,59^ Six samples were excluded from the analysis, one was determined to not be *Salmonella*, one appeared to be comprised of multiple genotypes, and four samples were on a long branch length and were conclused to be contaminated. Paired end reads for the remaining 464 samples were mapped to the *S*. Typhi CT18 reference genome (accession number: AL513382)^60^ using the RedDog mapping pipeline (v1beta.10b, available at http://githib.com/katholt/reddog). RedDog uses Bowtie2 v2.2.9^61^ to map all raw reads to the CT18 reference genome and then uses SAMtools v1.3.1^62^ to identify high quality SNP calls. SNPs that did not meet predefined criteria (a minimal phred quality score of 30 and depth coverage of 5 were filtered out).^63^ A failed mapping sequence was defined as when <50% of total reads mapped to the reference genome. 2 isolates were excluded from additional analysis after mapping failed, due to depth coverage of less than 10 (as per the RedDog pipeline default). A concatenation of core SNPs that were present in >95% of all genomes was generated and filtered to exclude all SNPs from phage regions or repetitive sequences in the genome reference CT18 as defined previously (Table S2).^60^ Gubbins (v2.3.2)^64^ was used to filter out SNPs in recombination regions. Finally, the alignment of 17,325 SNPs from mapping of the remaining 462 isolates was utilized for phylogenetic analysis. Resultant BAM files for all isolates from RedDog mapping were used to determine previously defined genotypes according to an extended genotyping framework using the GenoTyphi pipeline^65^ (available: https://github.com/katholt/genotyphi).

### Phylogenetic analysis

RAxML (v8.2.9)^26^ was used to infer maximum likelihood (ML) phylogenetic trees from the final chromosomal SNP alignment, with a generalized time-reversible model, a gamma distribution to model site-specific rate variation (the GTR+ Γ substitution model; GTRGAMMA in RAxML), and 100 bootstrap pseudo-replicates to assess branch support. *Salmonella* Paratyphi A AKU1_12601 (accession no: FM200053)^66^ was used as an outgroup. The resultant trees were visualized using Interactive Tree of Life (iTOL)^67^ and the ggtree package in R.^68^ An interactive visualisation of this phylogeny and associated metadata can be found in Microreact (https://microreact.org/project/hzELvWqY3UCvsyAw892fnd-origins-of-h58-s-typhi).,^69^

### Characterisation of AMR associated genes and mobile elements

SRST2 (v0.2.0) ^70^ was used to detect AMR genes and plasmid replicons using the ARGannot^71^ and PlasmidFinder^72^ databases, respectively. Mutations in the *gyrA* and *parC* genes, as well as the R717Q mutation in *acrB*, were detected using Mykrobe v0.10.0.^73^

### Bayesian phylogenetic analysis of H58 and nearest neighbours

Our estimation of the temporal signal of our H58 and nearest neighbour data exhibited a strong correlation between the sampling dates and the root-to-tip distances, with a positive value for the slope and an R^2^ value of 0.4743 (Supplementary Figure 2). Additionally, the randomly reassigned sampling time of sequences 20 times to generate the mean rates indicated that there was no overlap between the 95% credible intervals of the mean rate of the real data set and that of the date randomization data (Supplementary Figure 3). To infer where and when the first H58 (genotype 4.3.1) organism emerged, we conducted Bayesian phylogenetic analyses on a subset (n=345) of H58 (genotype 4.3.1) from our dataset and from published literature isolated between 1980 and 2000.^14,30,33^ This analysis of 345 *S*. Typhi isolates was conducted in the BEAST v1.8.4.^74^ The temporal signal of the data was checked initially. The maximum likelihood tree, constructed using the GTR+ Γ substitution model and GTRGAMMA, was subjected to TempEst v1.5 to test the best fit of linear regression between sampling dates and their root-to-tip genetic distances, using default TempEst parameters.^75^ To further test temporal signal, the TipDatingBeast R package was used to randomly reassign the sampling dates of sequences 20 times to create date-randomized data sets. BEAST analyses were conducted for these randomized data sets and the mean rates were compared between runs. The data had sufficient temporal signal if the 95% credible interval of mean rates of the date-randomized datasets did not overlap with that of the original sampling dataset.^76,77^

An automatic model selection program (ModelFinder)^78^ was implemented through IQ-TREE^79^ and run on the non-recombinant SNP alignment (724 variable sites) to select the best-fit sequence evolution model for BEAST analysis. ModelFinder showed that GTR had the lowest Bayesian Information Criteria (BIC) score and thus it was chosen as the best-fit substitution model.

As part of the BEAST analysis, six different model combinations were run for six combinations, and the final analysis was conducted using the best fitting model The path sampling and stepping-stone sampling approaches were applied to compare the log marginal likelihoods of the different runs.^36,80^ The GTR+Γ4 with strict clock and Bayesian skyline was identified as the best-fit model for running BEAST. Finally, BEAST was run three independent times using the best-fit model, using a Bayesian Markov chain Monte Carlo (MCMC) parameter-fitting approach (generated 10^7^ chains and sampled every 1000 iterations). The log files after three runs were combined using LogCombiner v1.8.3^81^ with a burn-in rate of 10%. The effective sample size (ESS) of all parameters was assessed by Tracer v1.8.3.^82^ If the ESS of any parameters was less than 200, we increased the MCMC chain length by 50% and reduced the sampling frequency accordingly.^36^ The trees were combined and summarized using LogCombiner v1.8.3 and TreeAnnotator v1.8.3.^74^

## Supplemental Figure Legends

**Figure S1**. A genetic comparison of IncH1 plasmid content of early H58 *S*. Typhi isolates. Zoomed in view of maximum likelihood phylogenetic tree including historical and published H58 isolates (n=206). Country of origin is indicated by the coloured circles at the tips of the tree. Gene content of an almost identical IncH1 plasmid is indicated by the colour to the right of the tree.

**Figure S2**. An estimation of temporal signal in H58 *S*. Typhi data.

Path-O-Gen/TempEst results (analysis included n=345 isolates using an alignment of 724 non-recombinant SNPs), demonstrating strong correlation between sampling dates and root-to-tip distance (R^2^ = 0.4743).

**Figure S3**. A TipDatingBeast estimation of temporal signal in H58 *S*. Typhi data. TipDatingBeast results (analysis incorporated n=345 isolates and an alignment of 724 non-recombinant SNPs), demonstrating no overlap between the original mean rates of mutation and mean rates of date randomization.

## Supplementary Tables

**Table S1**. Organism-level data and metadata for historical UKHSA *S*. Typhi isolates.

**Table S2**. Organism-level data and metadata for published H58 and nearest neighbours *S*. Typhi isolates.

**Acknowledgements**

Marie Anne Chattaway is affiliated to the National Institute for Health Research Health Protection Research Unit (NIHR HPRU) in Genomics and Enabling Data at University of Warwick in partnership with the UK Health Security Agency (UKHSA), in collaboration with University or Cambridge and Oxford. Marie Anne Chattaway is based at UKHSA. The views expressed are those of the author(s) and not necessarily those of the NIHR, the Department of Health and Social Care or the UK Health Security Agency

## Author Contributions

Conceptualization: SB

Data Curation: MEC, TNTN, TDHN, SN, MC

Formal Analysis: MEC, TNTN, ZAD, PTD

Funding Acquisition: SB

Investigation: MEC, TNTN, TDHN, ZAD, PTD, EM,

Resources: SN, MC

Project Administration: SB, MC Supervision: SB

Writing – Original Draft Preparation: MEC, SB

Writing – Review & Editing: MEC, TNTN, TDHN, ZAD, PTD, EM, SN, MC, SB

