## Supplementary figures and images for "The origins of haplotype 58 (H58) *Salmonella enterica* serovar Typhi"

### Supp Figure 1

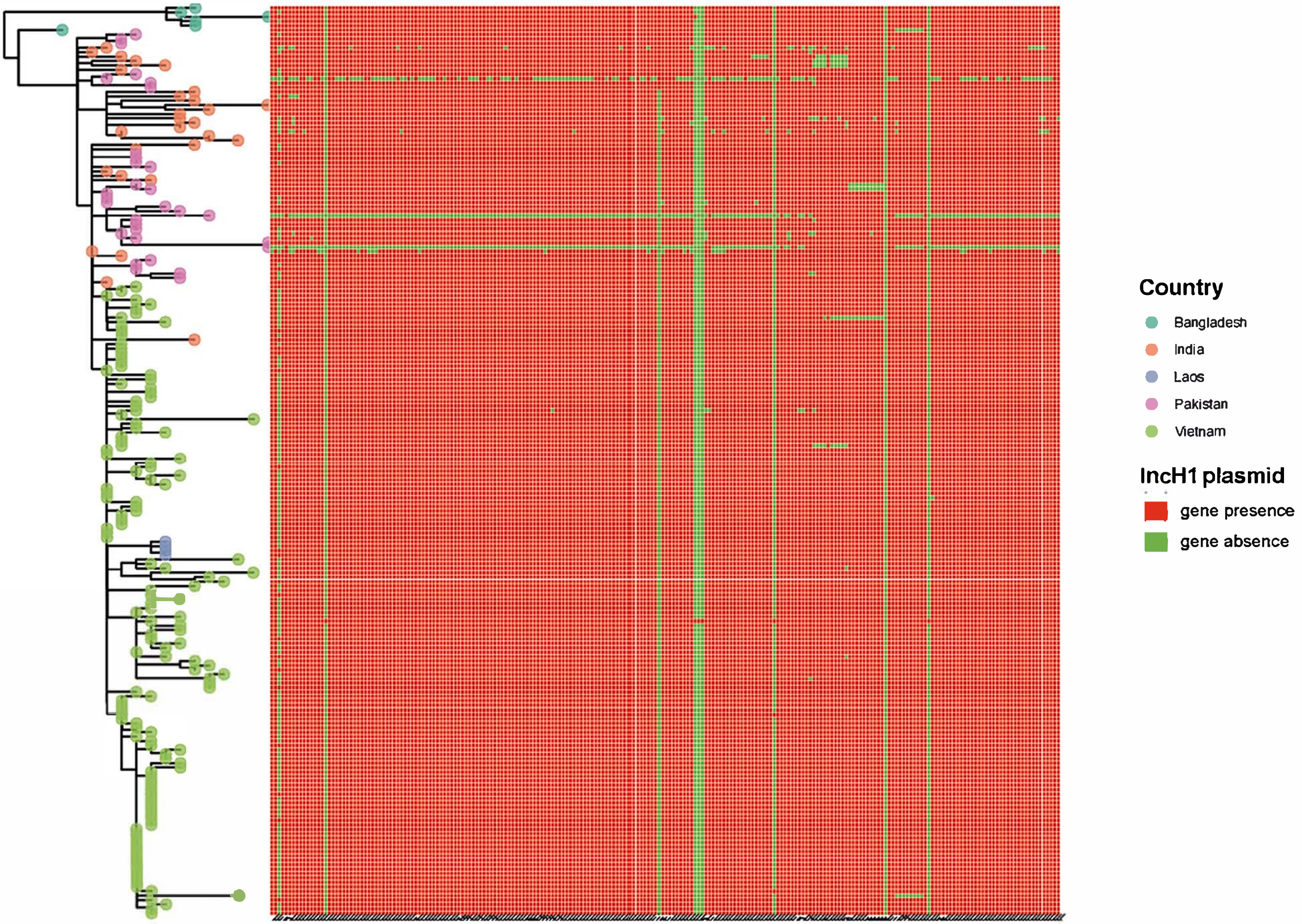

### Supp Figure 2

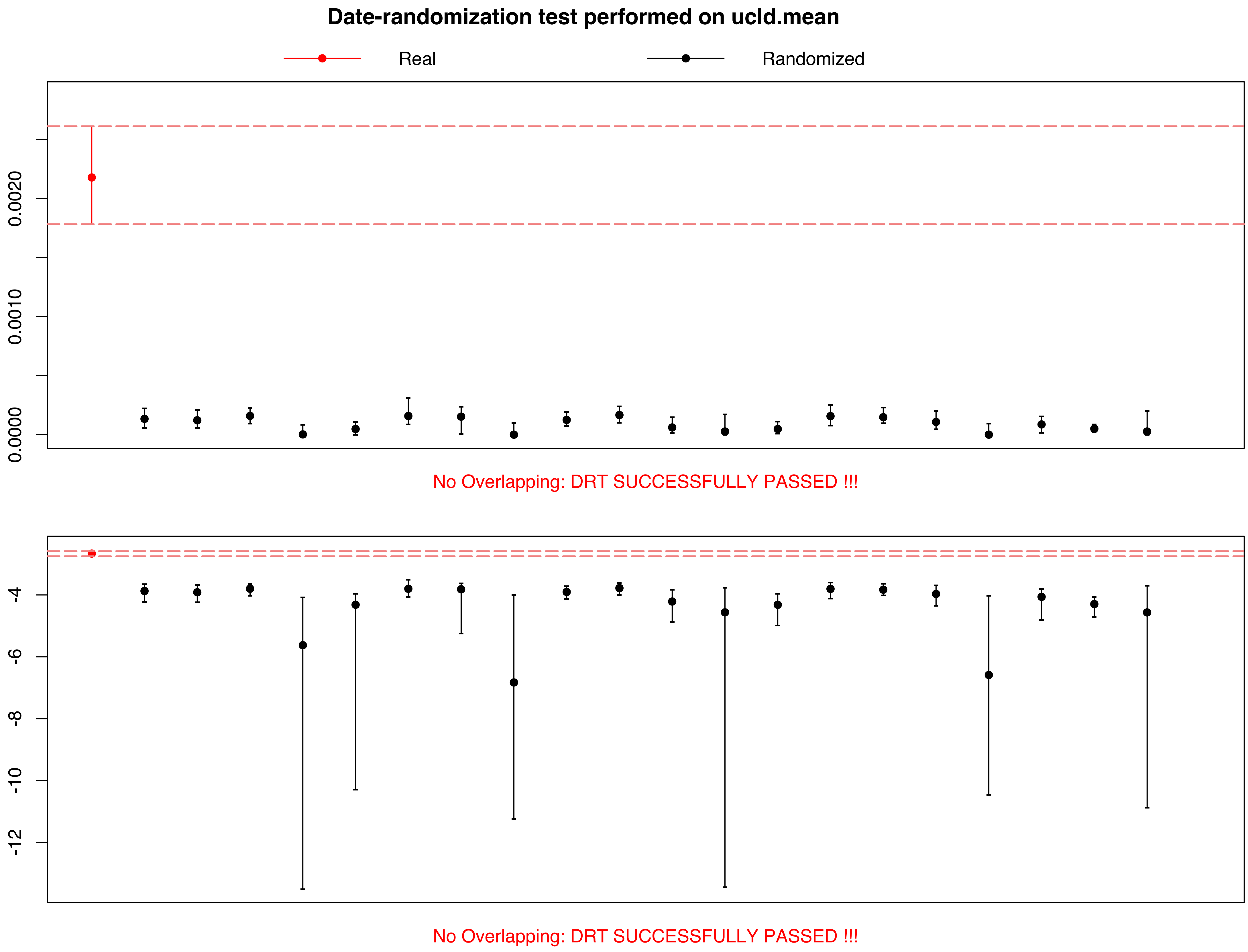
