## Supplementary material for "The origins of haplotype 58 (H58) *Salmonella enterica* serovar Typhi": Supp Figure 3

Best-fitting root

Function: heuristic residual mean squared

| Dated Tips | |
|---|---|
| Date range | 17 |
| Slope (rate) | 1.7153E-4 |
| X-Intercept (TMRCA) | 1949.1321 |
| Correlation Coefficient | 0.6887 |
| R squared | 0.4743 |
| Residual Mean Squared | 2.0781E-7 |

Sample Dates | Tree | Root-to-tip | Residuals | Node density

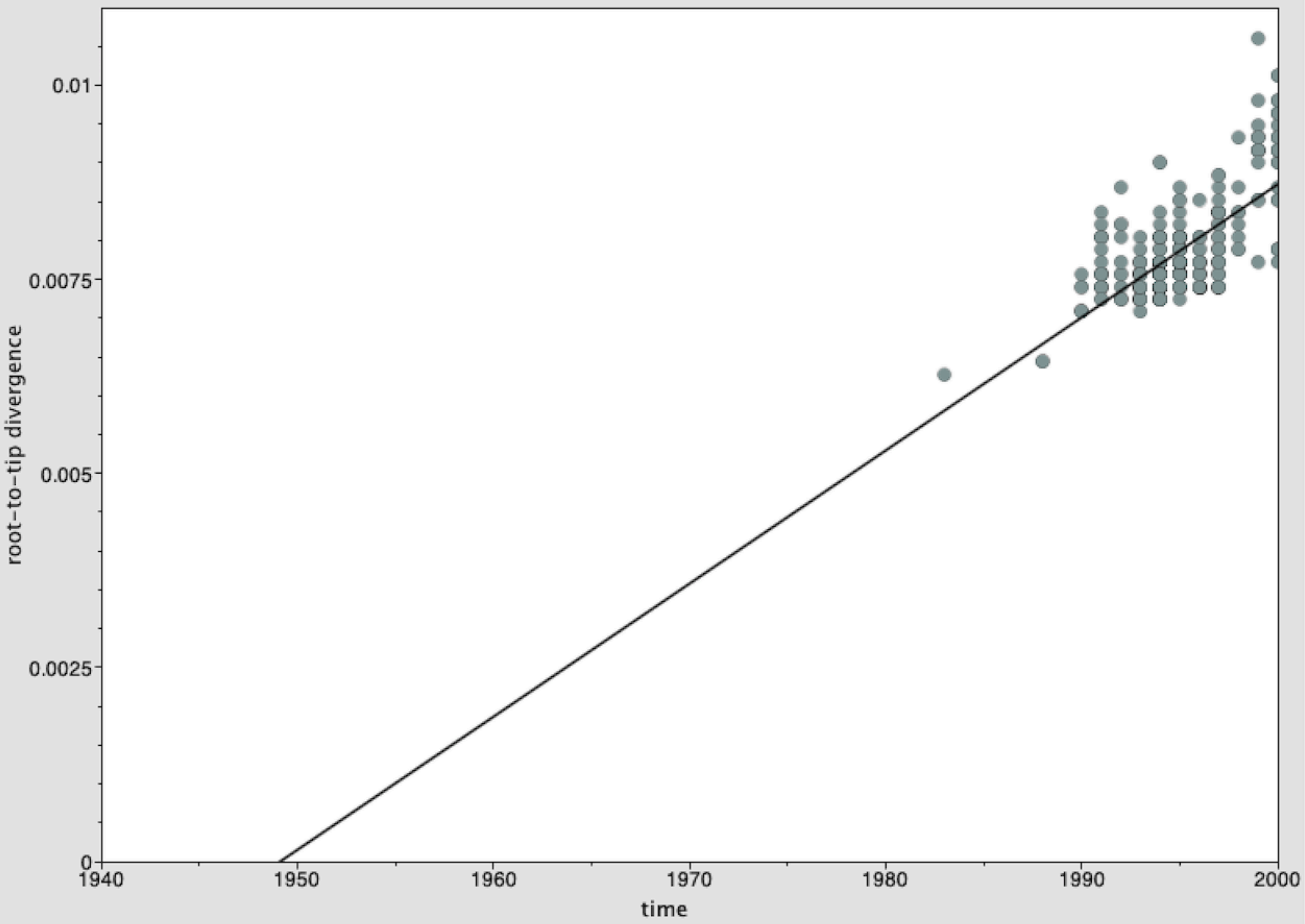


Show ancestor traces

Tree loaded, 345 taxa, dated tips with range 17
